# Variation in inbreeding depression within and among *Caenorhabditis* species

**DOI:** 10.1101/2025.06.04.657873

**Authors:** Matthew V. Rockman, Max R. Bernstein, Derin Çağlar, M. Victoria Cattani, Audrey S. Chang, Taniya Kaur, Luke M. Noble, Annalise B. Paaby

**Author notes:** Author for correspondence Matthew Rockman, Department of Biology, New York University, 100 Washington Square East, New York, NY 10003.

## Abstract

Outbreeding populations harbor large numbers of recessive deleterious alleles that reduce the fitness of inbred individuals, and this inbreeding depression potentially shapes the evolution of mating systems, acting as a counterweight to the inherent selective advantage of self-fertilization. The population biological factors that influence inbreeding depression are numerous and often difficult to disentangle. We investigated the utility of obligately-outcrossing (gonochoristic) *Caenorhabditis* nematodes as models for inbreeding depression. By systematically inbreeding lines from ten populations and tracking line extinction, we found that inbreeding depression is universal but highly variable among species and populations. Inbreeding depression was detected across the life cycle, from mating to embryo production to embryonic viability and larval growth, and reciprocal crosses implicated female-biased effects. In most cases, the surviving inbred lines have dramatically reduced fitness, but the variance among inbred lines is substantial and compatible with the idea that inbreeding depression need not be an obstacle to the evolution of selfing in these worms. Populations of some species, including *Caenorhabditis becei*, exhibited modest inbreeding depression and could be tractable laboratory models for gonochoristic *Caenorhabditis*.

## INTRODUCTION

The constant generation of damaging mutations saddles populations with a genetic load. This load consists of an expressed component, manifest as the reduction of fitness of an individual drawn from the population relative to that of a mutation-free individual, and a concealed component, due to recessive mutations, that manifests as inbreeding depression, a reduction of fitness in inbred relative to outbred individuals (Morton *et al*. 1956; Lynch and Walsh 1998; Charlesworth and Willis 2009; Hedrick and Garcia-Dorado 2016; Dussex *et al*. 2023).

Many aspects of population biology influence the distribution of expressed and concealed load. In small populations, deleterious alleles drift from their expected frequencies, resulting in higher expressed load (drift load) than in larger populations (Kimura *et al*. 1963). While weakly deleterious mutations can drift all the way to fixation in small populations, these populations have more efficient selection against recessive deleterious variants, as these variants find themselves homozygous more often in small than in large populations. Overall, theory predicts that large populations will harbor more concealed load and less expressed load than smaller populations (Bataillon and Kirkpatrick 2000; Robinson *et al*. 2023). At the same time, population structure, particularly matings among close relatives, can also reduce concealed load. Inbreeding purges recessive deleterious variation by increasing the selectable phenotypic variance, both by genetic drift and by systematic effects on genotype frequencies (Glemin 2003).

Efforts to study broad patterns of relationship among expressed and concealed load, population history, and genetic diversity have been hampered by the difficulties in measuring the relevant quantities in populations that differ in these respects but are otherwise similar in their genetics and life history. Here, we focus on a set of related species that are extremely similar in basic biological parameters. *Caenorhabditis* nematodes exhibit exceptional morphological and molecular conservatism, right down to cell lineage (Zhao *et al*. 2008; Memar *et al*. 2019), developmental gene expression (Levin *et al*. 2012), gene function (Verster *et al*. 2014), and chromosome organization and recombination landscape (Stein *et al*. 2003; Ross *et al*. 2011; Teterina *et al*. 2020; Noble *et al*. 2021; Sun *et al*. 2022; Teterina *et al*. 2023), and they are well matched in factors such as offspring number and body size.

The best studied *Caenorhabditis* species have androdioecious mating systems, and they occur in nature primarily as highly inbred self-fertile hermaphrodites, incapable of mating with one another. Rare males provide opportunities for outcrossing. The majority of *Caenorhabditis* species, conversely, are obligate outcrossers with separate males and females (gonochorism). The contrast in mating systems is associated with profound differences in the magnitude of inbreeding depression, which is effectively absent in the selfing species and extreme in the best-studied outcrossers, and genetic diversity, which is depauperate in selfers and preposterous in outcrossers (Dolgin *et al*. 2007; Dey *et al*. 2013; Cutter *et al*. 2019; Braendle and Paaby 2024). Efforts to study gonochoristic species have been severely hampered by the difficulty of making inbred lines; lines propagated by full-sib mating rapidly suffer extinction (Dolgin *et al*. 2007; Barriere *et al*. 2009; Fierst *et al*. 2015; Teterina *et al*. 2020).

As inbreeding depression can act as a barrier to mating-system evolution (Lloyd 1979), the levels of segregating recessive deleterious variation in extant male/female *Caenorhabditis* species may be an effective barrier to transitions from gonochorism to selfing. The clade has given rise to extant species with androdioecious mating systems three times independently (Kiontke *et al*. 2011), and the genetic requirements for such mating system transitions are minimal: *C. remanei* can be converted from gonochorist to selfer by reducing the activity of just two genes (Baldi *et al*. 2009).

The phenotypic manifestations of inbreeding depression depend on the nature of the underlying recessive genetic variants, and these in turn are influenced by the interaction of population biology, developmental biology, and selection. Genes acting in males and females experience each of these factors differently, potentially generating sex-biased inbreeding depression (reviewed in Ebel and Phillips 2016). Maternal genetic effects may play a special role: these are sex-limited in their expression and so are expected to harbor twice as much variation at mutation-selection equilibrium as zygotic-effect genes (Wade 1998). They also superimpose among-family selection on top of selection acting among individuals, and they induce positive frequency-dependence when they genetically interact with offspring genotypes, which is compounded by the elevated parent-offspring genetic correlations under inbreeding (Wolf and Wade 2016).

We experimentally estimated loads of concealed and exposed deleterious variation in isolates of several gonochoristic *Caenorhabditis* species. We set out to address four questions. First, to what extent do gonochoristic species vary in their loads of concealed and expressed genetic load? Second, are there gonochoristic species in which the concealed load is sufficiently low that they can be inbred to establish a convenient experimental model species for gonochoristic *Caenorhabditis*? Third, is the concealed load in gonochoristic species large enough to act as an obstacle to evolutionary transitions to selfing? And fourth, does inbreeding depression manifest differently across life stages and sexes?

## METHODS

### Establishment of experimental populations

We experimentally inbred eleven isolates of gonochoristic *Caenorhabditis* nematodes and measured the change in cross success with increasing levels of inbreeding. Two isolates of *C. remanei*, EM464 from Brooklyn, New York, and PB219 from Dayton, Ohio, are widely studied exemplars of this cosmopolitan, genetically diverse species and are included as positive controls for the presence of inbreeding load (Dolgin *et al*. 2007; Barriere *et al*. 2009; Fierst *et al*. 2015; Ebel and Phillips 2016; Adams *et al*. 2022). At the other end of the spectrum, we examined CB4108, a *fog-2(q71)* mutant strain of *C. elegans* (Schedl and Kimble 1988). This completely inbred derivative of the androdioecious lab strain, N2, from Bristol, England, is defective in hermaphrodite self-sperm production and therefore behaves as a gonochorist. This strain is expected to exhibit no inbreeding depression and is included as a negative control. EM464, PB219, and CB4108 were acquired from the *Caenorhabditis* Genetics Center. The remaining eight experimental populations are two strains from each of four gonochoristic species from across the Elegans supergroup: *C. panamensis* and *C. becei* from Barro Colorado Island, Panama (Sloat *et al*. 2022), *C. kamaaina* from Kaua’i, Hawaii (Félix *et al*. 2014), and *C. remanei* from Okinawa, Japan. We established each population from wild-collected animals via isofemale bottleneck, which isogenizes mitochondrial genotype in each population, followed by expansion to large population sizes (~10^5^) over several generations prior to cryopreservation of these founder stocks.

### Experiment 1: Estimating load parameters by serial sib mating

We thawed each strain and established large populations on 10cm NGM-agarose plates seeded with OP50-1 *E. coli* bacteria as a food source. Each population was cleaned of any contaminants by bleaching, preserving population size. Worms went through approximately eight overlapping generations under these conditions prior to the start of inbreeding. We then initiated 108 independent lines of each wild strain for inbreeding by serial sib-mating and 120 lines for *C. elegans fog-2* mutants (1200 experiments in total). Crosses involved pairing a single L4 (juvenile) male and a single L4 female on a 6cm NGM-agarose plate seeded with 50μl of OP50-1. All crosses were performed at room temperature and were carried out by six people, each starting simultaneously with 200 anonymized lines in a balanced randomized block design. Each cross yielded L4s of each sex to establish a new cross, or it failed to do so and was scored a line extinction. The primary phenotype scored is thus a binary outcome: does a pair of worms, conditional on having reached the L4 stage, succeed or fail in producing at least one L4 offspring of each sex. In addition to this basic cross success phenotype, we analyzed three subphenotypes: *Copulation*, the probability that a pair of worms successfully copulates, as demonstrated by the presence of a copulatory plug on the female; *Fertility*, the probability that a copulation results in laying of embryos; and *Growth*, the probability that some of the embryos develop to the L4 stage. Inbreeding was stopped in the 21^st^ generation. Surviving lines were cryopreserved and are available for further analysis.

Morton, Crow, and Muller (1956) modeled the probability of an individual’s survival under inbreeding as a function of several factors, including the effects of deleterious variants segregating at Hardy-Weinberg Equilibrium, the effects of environmental factors, and the effects of recessive deleterious variants homozygosed by inbreeding. Assuming that the effects of each locus and environmental factor are small and independent, *S*, the probability of survival, takes on a simple form, *S = e*^*–A – BF*^. *F* is the inbreeding coefficient, the probability that alleles in an individual are identical by descent. In a regression of −log *S* on *F*, the intercept *A* represents the effects of environmental factors, including those that depend on fixed load, and the effects of segregating load in the absence of inbreeding. *B* represents the change in survival probability as a function of inbreeding.

We modified this approach to model *R*, the probability of a successful reproduction event. Because reproduction requires that each of two worms succeeds, the coefficients *A*_*R*_ and *B*_*R*_ are approximately twice the magnitude of Morton *et al*.’s *A* and *B* for survival probability, as shown in File S1. We regressed the negative log of the observed proportion of successful crosses on strain and *F* within strain, iteratively reweighting observations according to the inverse of the expected error variance, as in Morton *et al*. (1956) (File S2). Defining the starting population as our reference, the first generation of each cross is between outbred (*F* = 0) individuals, yielding outbred embryos (*F* = 0).

Crosses in the next generation therefore also represent *F* = 0 crosses. Their offspring are inbred (*F* = 0.25), and sequential generations have inbreeding coefficients that increment each generation according to the recursion *F*_*t*_ = 0.25 * (1 + 2*F*_*t*-1_ + *F*_*t*-2_) (Lynch and Walsh (1998), eq. 10.5b).

The genetic nonindependence of sequential observations results in a downward bias in the estimates of the standard errors of the slopes in our regressions (Lynch 1988), but this bias is small in magnitude because of the large number of lines assayed per strain and the very small additive genetic variance for *R* in the founding populations. We verified, using Lynch’s *J* statistic, that our results are robust to this concern (File S2).

Some analyses involve observations where there are zero successes (i.e., for a particular strain at a particular *F*, no crosses succeeded), which results in a log(0) situation. We excluded these observations from the analysis, but that exclusion introduces a bias because the probability of zero successes increases as sample size decreases, even for constant probability of failure. We therefore used simulations with constant probability of failure to generate null distributions for statistical tests in the presence of zero-success observations (File S2). For each strain, we simulated the *t*-statistic for Strain:*F* terms under the null hypothesis *B* = 0. We fixed the probability of success in the simulations as the observed global mean for the dataset. The simulated *t* distributions were slightly shifted toward lower values, as expected. But the shifts were very small, <0.15 for each strain, and do not affect any conclusions.

### Experiment 2: Relative fitness of outbred populations and inbred lines

We estimated fitness from reproductive schedules for six females from each of nineteen strains: five outbred founders and two to three inbred lines derived from each. The inbred lines were selected at random from the collection of inbred lines from experiment 1.

We thawed each strain and maintained them on food at large population sizes for several generations prior to the start of the experiment. We isolated L4 animals of each sex and left them to mature. We then paired an adult male and an adult female for five hours, after which we removed the male. The female was left to lay embryos until six hours from the initial pairing and was subsequently transferred to a new plate every eight hours until it had stopped laying embryos. After 60-80 hours of development, we counted the adult progeny on each plate (*i*.*e*., dead embryos and arrested larvae were not counted). The experiment was performed at room temperature in a single block by a single investigator, blinded to strain identity, and the order of the 120 females was randomized.

Seven females were censored from analysis because they failed to mate or they crawled onto the edge of the plate and desiccated before the conclusion of egg laying. For the remaining 113 females, we calculated the total number of progeny that developed to adulthood and we estimated relative fitness using the Lotka equation for age-structured populations, as in Dolgin *et al*. (2007). The population growth parameter *r* was chosen to set the mean fitness of the outbred founder females to 1, separately for each founder (*i*.*e*., comparisons are between founders and their descendant inbred lines, not among founders or unrelated inbred lines). Note that fitness measured in this way incorporates the reproductive schedule starting from adulthood; development time to reproductive maturity for the assayed females was not measured but likely contributes to fitness. Relative fitness is correlated with number of offspring (R^2^ = 0.93) but better captures the proliferative ability of each animal in the case of overlapping generations.

### Experiment 3: Tests for complementation and maternal effects

We used scanner-based population growth-rate assays (Tintori *et al*. 2024) to estimate fitness complementation and maternal effects for two inbred lines of *C. panamensis*, QG1819 and QG1892, both derived from founder isofemale line QG702.

The strains were thawed and raised at 25^°^ C on NGMA plates seeded with OP50-1. Populations were then bleached and the resulting L4s picked to establish four types of cross: QG1819xQG1819, QG1892xQG1892, QG1819xQG1892, and QG1892xQG1819. When the progeny of these crosses had developed to L4, they were separated by sex and allowed to mature to adulthood overnight. The resulting first-day adults were then picked to 3.5 cm assay plates containing NGMA with 10µg/ml nystatin and 50µg/ml streptomycin. Assay plates were seeded with 200 µl of OP50-1, concentrated 10X from an overnight culture. Six types of plate were set up: self crosses of QG1819 and QG1892, reciprocal crosses between QG1819 and QG1892, and self crosses of each class of F_1_, those with QG1819 mothers and those with QG1892 mothers. Each assay plate received two males and two females, with ten replicate plates per cross type.

Plates were randomized across acrylic plate-holders in four manually refocused Epson V800 and V850 scanners, all inside a 25 °C incubator. Scanners recorded 1200-dpi images once per hour for 8 days. Images were processed and statistics derived following the *popscan* pipeline (Tintori *et al*. 2025). We compared log(hours to resource exhaustion) among strains using *t*-tests.

## RESULTS

We set up crosses between single male and female worms and scored whether the worms mated, whether the female laid embryos, and whether those embryos hatched and developed to the L4 stage. If the cross produced L4 progeny, one pair of male and female worms were randomly selected for the next generation; if the cross failed we recorded the step at which it failed: copulation, fertility, or growth. Of 1200 such experiments, 157 (13%) successfully reproduced through 20 consecutive generations of full-sib matings; the remaining lines suffered extinction during the experiment (Figure 1). The distribution of extinctions across the resulting 11,892 crosses revealed the patterns of expressed and concealed genetic load in 11 strains of gonochoristic *Caenorhabditis* nematode.

**Figure 1.**
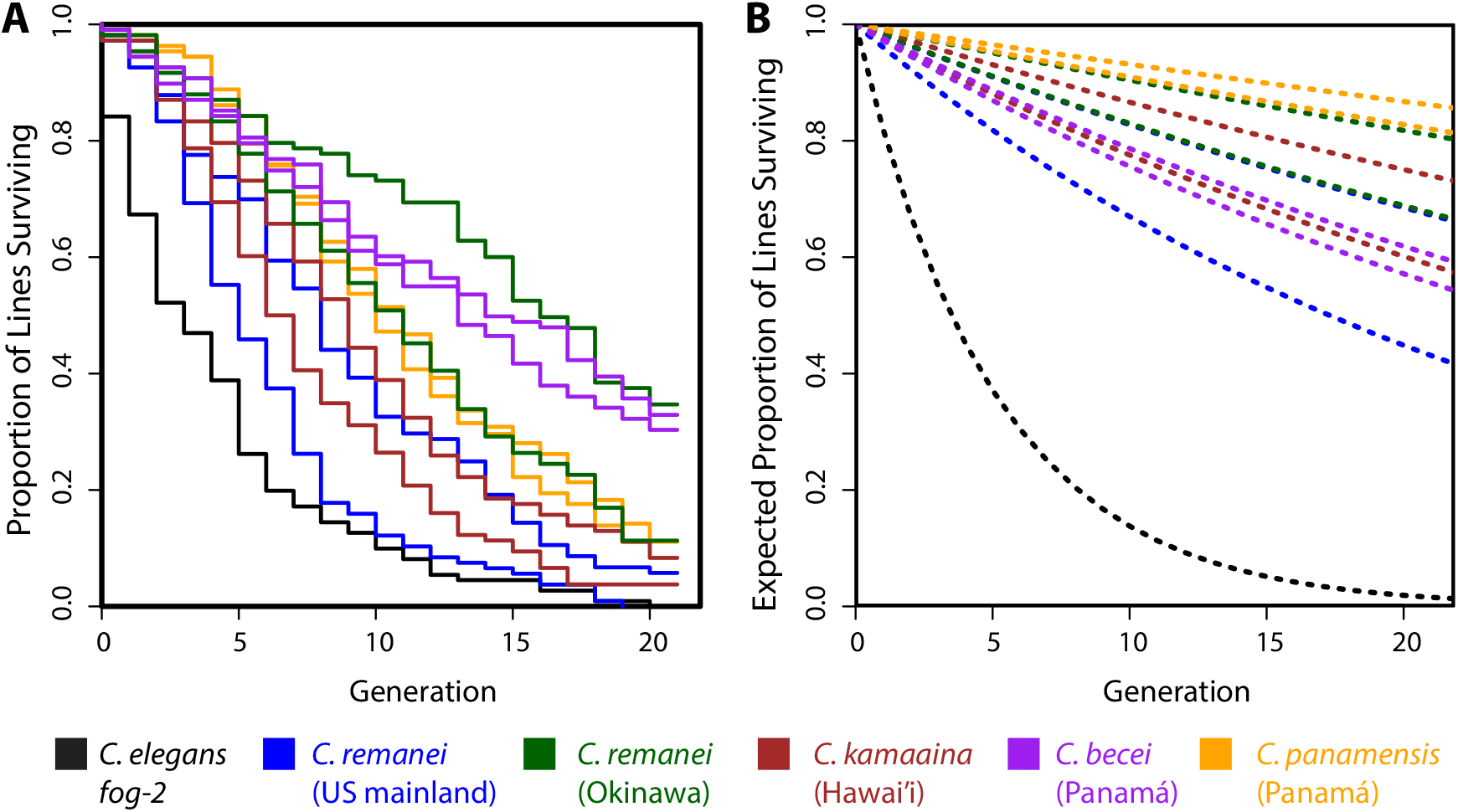
A. During the course of 20 generations of serial full-sib mating, the majority of lines for each of 11 *Caenorhabditis* isolates went extinct. The two lines for each color represent the two different founding isolates for each wild population. **B**. In the absence of inbreeding depression, the expected survival curves follow exponential decay with rate –*e*^*A*^, reflecting a constant probability of cross failure. The difference between panels A and B demonstrates the extent of concealed genetic load (*B*_*R*_), which is significant for every strain except the control strain, *C. elegans fog-2*. This strain began the experiment fully inbred, and its steep die-off reflects its high but constant rate of cross failure. See Table 1 for estimates, errors, and *p*-values.

Adapting the design of Morton *et al*. (1956), we used the regression of the logarithm of the probability of cross success on the inbreeding coefficient *F* to estimate the load for each isolate (Table 1). Every wild isolate showed significant concealed load, manifesting as an increase in extinction probability with increasing inbreeding (*i*.*e*., *B*_*R*_ > 0, *p* < 0.05). The negative control strain, *C. elegans fog-2*, which was completely inbred from the start, showed the expected absence of inbreeding depression. The severity of inbreeding depression varied considerably among species, and, in some cases, among isolates within species (Table 1).

**Table 1.** Estimates of expressed and concealed load. Nominally significant estimates are bolded.

| Species | Strain | Locality | $\hat{A}_R \pm \text{SE}$ | $\hat{B}_R \pm \text{SE}$ | $p_A$ | $p_B$ |
| --- | --- | --- | --- | --- | --- | --- |
| *C. elegans* | *fog-2* | NA | **$0.198 \pm 0.031$** | $0.051 \pm 0.068$ | $<10^{-8}$ | 0.452 |
| *C. remanei* | EM464 | New York | $0.019 \pm 0.010$ | **$0.163 \pm 0.026$** | 0.058 | $<10^{-8}$ |
| *C. remanei* | PB219 | Ohio | **$0.040 \pm 0.015$** | **$0.299 \pm 0.048$** | 0.007 | $<10^{-8}$ |
| *C. remanei* | QG548 | Okinawa | $0.010 \pm 0.007$ | **$0.120 \pm 0.018$** | 0.164 | $<10^{-9}$ |
| *C. remanei* | QG549 | Okinawa | **$0.019 \pm 0.009$** | **$0.040 \pm 0.014$** | 0.040 | 0.006 |
| *C. kamaaina* | QG122 | Kaua'i | $0.014 \pm 0.009$ | **$0.145 \pm 0.023$** | 0.097 | $<10^{-9}$ |
| *C. kamaaina* | QG123 | Kaua'i | **$0.025 \pm 0.012$** | **$0.204 \pm 0.033$** | 0.028 | $<10^{-8}$ |
| *C. panamensis* | QG702 | Panama | $0.009 \pm 0.007$ | **$0.124 \pm 0.019$** | 0.178 | $<10^{-9}$ |
| *C. panamensis* | QG703 | Panama | $0.007 \pm 0.006$ | **$0.119 \pm 0.017$** | 0.247 | $<10^{-10}$ |
| *C. becei* | QG704 | Panama | **$0.028 \pm 0.011$** | **$0.040 \pm 0.018$** | 0.011 | 0.022 |
| *C. becei* | QG711 | Panama | **$0.024 \pm 0.010$** | **$0.040 \pm 0.016$** | 0.019 | 0.015 |

Several isolates showed significant expressed load in the absence of inbreeding (i.e., *A*_*R*_ > 0, *p* < 0.05). These values, which reflect the fixed and segregating load carried by each isolate, were small in magnitude (Table 1). The exception is the *C. elegans fog-2* control, for which *A* was approximately 0.2; this value corresponds to a cross failure rate of about 18%.

Crosses failed at three different steps in the reproductive process: copulation, fertility, and growth. Analyzing these steps one by one, we found that the expressed load (*A*) was in all cases attributable to the copulation step; conditional on successful copulation, we saw no significant expressed load for fertility and growth (Table S2). In contrast, most strains showed significant inbreeding load (*B*) for each step of the process. That is, we observed inbreeding depression for copulation success, and conditional on copulation success we observed inbreeding depression for fertility, and conditional on fertility we observed inbreeding depression for embryonic and larval growth. Several exceptions may point to potential differences among strains in the characteristics of their recessive deleterious alleles. For example, *C. becei* QG704 shows significant load for fertility but not growth, while *C. becei* QG711 shows the opposite pattern.

Isolates that show low inbreeding depression by our binary reproductive success metric may nevertheless suffer considerable quantitative reductions in fitness. To assay fitness quantitatively, we selected five outbred founder strains (EM464, QG549, QG123, QG702, and QG704) and two to three random inbred lines derived from each. For these 19 strains, we estimated the fitness of the inbred derivatives relative to their outbred ancestors by measuring the reproductive schedules and number of offspring that developed to adulthood.

In most cases, inbred lines exhibited significantly reduced fitness relative to their outbred ancestor (Figure 2). For *C. remanei* EM464, the fittest of the three descendant inbred lines had a relative fitness of only 0.135. Each of the other founders produced a mix of severely damaged (relative fitness < 0.5) and reasonably fit inbred descendants, and in one case, a line derived from *C. becei* QG704, inbred descendants had higher average relative fitness than their outbred ancestors. For 3 of 5 founders, the inbred lines from each exhibited significant heterogeneity in fitness (*F* tests for effect of strain, *p* < 0.05).

**Figure 2.**
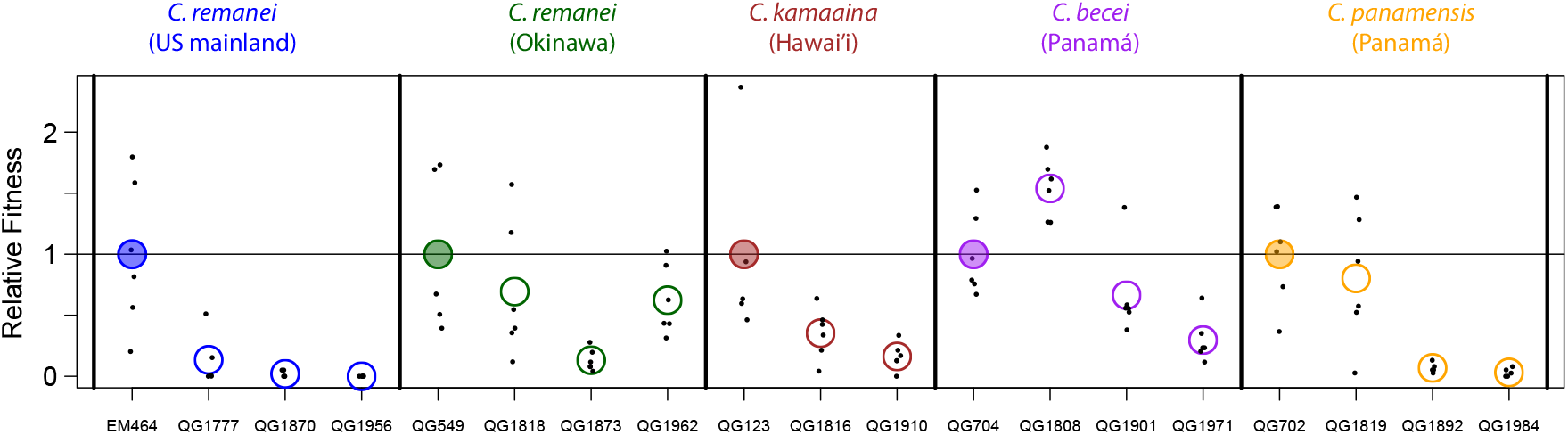
Inbred lines have reduced and heterogeneous fitness relative to their ancestor. The *y*-axis shows the relative fitness of the five ancestors and several of their inbred descendants. Each ancestor’s mean relative fitness (filled circles) is set to 1. Each point is a single mated female, and the circles represent strain means.

If inbred lines derived from a single founder population have fixed different haplotypes, crosses between them should result in complementation of recessive alleles and a restoration of fitness. We tested for complementation using two *C. panamensis* lines derived from founder QG702. The two inbred lines, QG1819 and QG1892, differ dramatically in relative fitness in our reproductive schedule assay (Figure 2), and so are likely to have fixed different haplotypes. We used flatbed scanner assays to estimate population fitness in these two lines, reciprocal crosses between them, and crosses between F_1_ animals from each of the reciprocal crosses (Figure 3). These assays recapitulated the large difference in fitness between the inbred lines (*t*-test, *p*<10^−7^), and crosses result in a dramatic increase in fitness relative to the less fit parent, consistent with complementation.

**Figure 3.**
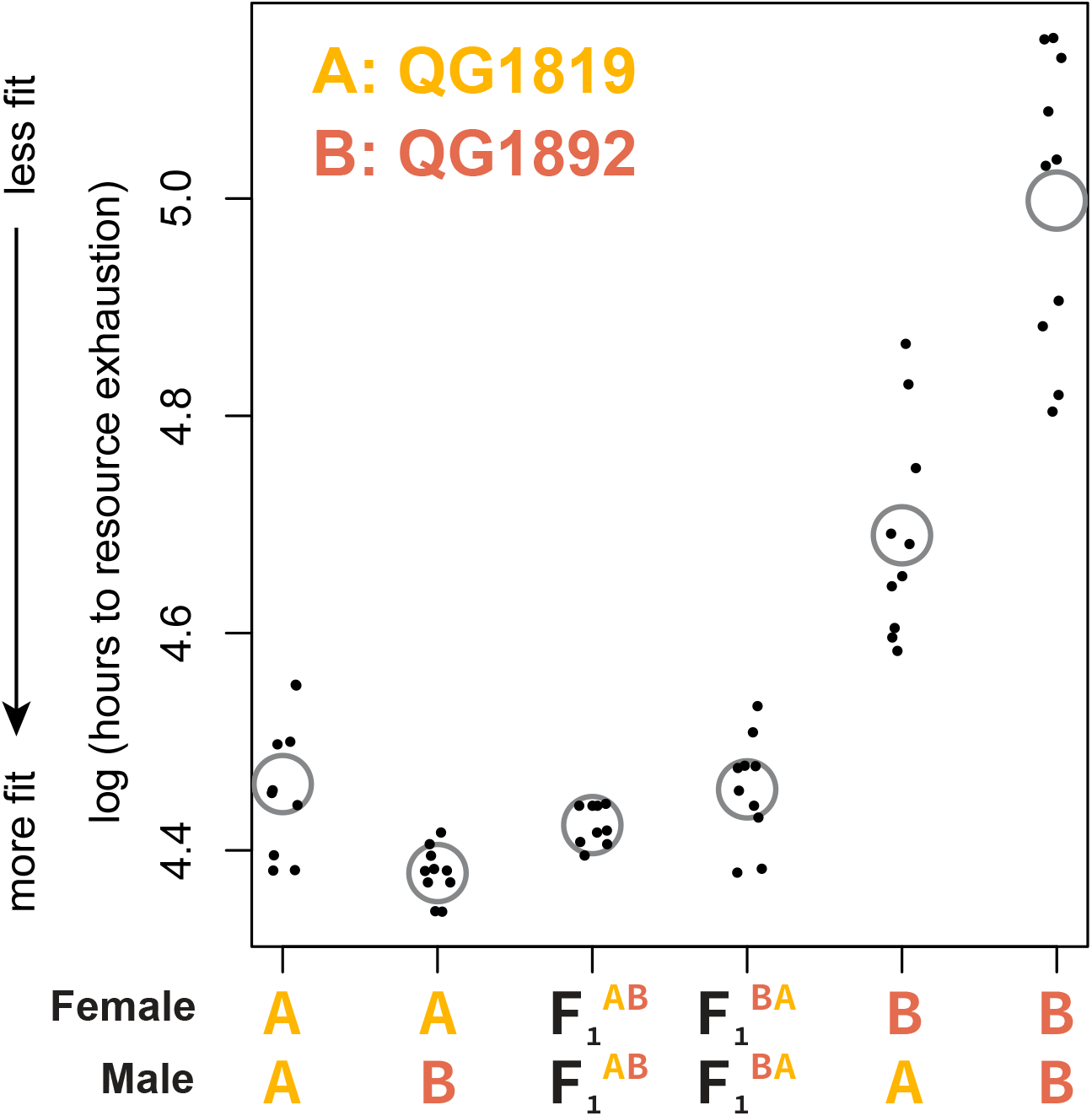
Crosses between related inbred lines of *C. panamensis* show complementation and maternal effects. The *y*-axis shows the log of the number of hours required for a population founded by two males and two females to exhaust a fixed quantity of resources; the population at that point typically includes thousands of grandchildren of the founders. Points lower on the *y*-axis represent populations with faster population growth and consequently greater fitness. Each point is a population and gray circles indicate means. The F_1_s are the product of matings between females of strain A and males of strain B (F_1_^AB^) or the reciprocal cross (F_1_^BA^).

Notably, reciprocal crosses between the inbred lines differ substantially (*p* <10^−5^), consistent with sex-specific inbreeding depression: fitness is much higher when the high-fitness inbred line is the female in the cross. This directional effect is reduced or absent when crosses start with the F_1_ worms from the reciprocal crosses (*p* = 0.08). Given our experimental design, this pattern implicates female-biased inbreeding depression that results in some mixture of reduced brood size or ovulation rate (features of the maternal phenotype) or reduced viability or developmental rate in the F_1_s (maternal-effect features of the pre-adult F_1_ phenotype). The frequencies of the two versions of the X chromosome also differ in reciprocal crosses and may contribute to phenotypic difference.

## DISCUSSION

We found that all tested gonochoristic *Caenorhabditis* isolates carry a significant burden of recessive deleterious alleles, with fitness declining as inbreeding increases. These results are congruent with findings from experimental analyses of *C. remanei* and *C. brenneri* (Dolgin *et al*. 2007; Barriere *et al*. 2009; Fierst *et al*. 2015; Ebel and Phillips 2016; Adams *et al*. 2022), and with the commonplace experience of every researcher working with gonochoristic *Caenorhabditis* strains.

The severity of the experimentally exposed inbreeding depression varies substantially among species and among isolates within species. In *C. remanei*, a widely distributed generalist species, we observe higher concealed load in two strains from North America than in two strains from Okinawa. This observation is consistent with the theoretical expectation that reduced population size, associated here with life on a small island, should facilitate purging of concealed load (Dussex *et al*. 2023). However, *C. kamaaina* isolates, from an island of similar size but greater remoteness (Kauai), has a very high concealed load. *C. kamaaina* is likely an ancient Hawaiian endemic, while Okinawan *C. remanei* may be a recent peripheral isolate; thus population age may be playing a key role in the architecture of recessive load. *C. becei* and *C. panamensis*, nearly indistinguishable species that live in sympatry in Panamá (Sloat *et al*. 2022), differ in their estimated loads by a factor of three. These disparate levels of concealed load reinforces the view that *Caenorhabditis* nematodes are a useful model for discovering the population-biological factors that shape recessive deleterious variation.

A key practical finding is that *C. becei* exhibits a manageably low concealed load, with a large number of its experimental lines (70/216) surviving 20 generations of sib mating. By comparison, only 10/216 lines of the North American *C. remanei*, the classic model gonochoristic *Caenorhabditis*, survived the process. In addition, some of the *C. becei* inbred lines retained high fitness. Given these observations, we are now developing tools for experimental studies of *C. becei*, to establish it as a tractable model for obligately sexual *Caenorhabditis* species and for genetic analysis of recessive deleterious alleles (Salome-Correa *et al*. 2025).

Some isolates, but not all, also have signatures of expressed genetic load prior to systematic inbreeding, perhaps as a symptom of their original derivation by isofemale bottleneck. By far the greatest level of expressed load is found in *C. elegans*. As expected, the *fog-2* mutant strain exhibits no significant inbreeding depression, because it is already fully homozygous. But it is profoundly deficient at mating, failing in 18% of pairings. Some fraction of this deficit may reflect the fixation of deleterious alleles in the ancestry of *C. elegans*, associated with the evolution of selfing or with the accumulation of deleterious alleles during long periods of evolution with little outcrossing and reduced effective population sizes (Cutter 2019). However, the step in our assay at which *C. elegans* fails – copulation – suggests that the expressed load in *C. elegans* may simply reflect relaxed selection on male function and mating ability in this androdioecious species, which almost always reproduces by self-fertilization in nature (Palopoli *et al*. 2008; Noble *et al*. 2015; Yin *et al*. 2018; Cutter *et al*. 2019).

In theory, the load of recessive deleterious alleles carried by a gonochoristic population presents an obstacle to mating-system evolution via invasion by selfing alleles (Lloyd 1979; Charlesworth and Charlesworth 1987). Although we observed significant decreases in the probability of successful reproduction in each of the ten gonochoristic isolates we examined, the magnitudes of these decreases were not large. Estimated cross success at *F* = 1 ranges from 71% in PB219 to 98% in QG549. Our experiments involved single-worm-pair matings, but in natural populations, typical brood sizes in the high hundreds would buffer the extinction risk of even dramatically lowered per-pair reproductive success probabilities. The worm-breeder’s frustration with experimental inbreeding may largely reflect our historic reliance on strict sib-mating designs.

If cross success were a comprehensive measure of individual fitness, inbreeding depression in these isolates is sufficiently small that an allele causing selfing should be able to invade. Because the transition to selfing in *Caenorhabditis* is genetically simple (Baldi *et al*. 2009), these data suggest that the true fitness costs to inbreeding may greatly exceed those measured by our assay sib-mating assay. We observed such costs in our analysis of reproductive schedules for fully inbred individuals (Figure 2). In many cases – in all three genotypes examined for the North American *C. remanei* – relative fitness was substantially lower than the canonical value required to prevent invasion of selfing (Lloyd 1979). In most species, however, we observed substantial among-line variation in the reduction in fitness, with some lines performing very well. This variance among individuals in the burden of recessive deleterious alleles creates the potential for facile transition to mixed-mating and ultimately selfing (Lande and Schemske 1985; Ebel and Phillips 2016; Brown and Kelly 2020). Such transitions would potentially be aided by progressive purging of the recessive load and benefits to selfers from mating assurance in patchy environments. Inbreeding depression may not be the whole explanation for the rarity of selfing species in *Caenorhabditis*.

Our experiments allow us to localize inbreeding depression in several different places in the life cycle. We found that, for most isolates, inbreeding load was detectable at the steps of mating, fertility, and embryonic and larval growth. In the one pair of *C. panamensis* inbred lines we examined in reciprocal crosses, we detected a strong female bias in the magnitude of inbreeding depression. These findings generalize the results of previous studies in *C. remanei* (Dolgin *et al*. 2007; Ebel and Phillips 2016).

Female-specific effects in our assay likely represent a mixture of direct and maternal-effect phenotypes. Anecdotally, each of the ten wild isolates we inbred produced at least one line that went extinct when the female laid a large number of embryos, all of which failed to hatch. This is a typical maternal-effect pattern, with no segregation within the brood. Maternal-effect genetic variation is abundant within and among *Caenorhabditis* species (Farhadifar *et al*. 2015; Ben-David *et al*. 2017; Farhadifar *et al*. 2020; Noble *et al*. 2021), as is variation in zygotic sensitivity to maternal genetic perturbations (Paaby *et al*. 2015; Torres Cleuren *et al*. 2019). While maternal-effect inbreeding depression in vertebrates putatively operates through changes in maternal care (*e*.*g*., Sin et al. 2021), the cytoplasmic basis of maternal genetic effects in *Caenorhabditis* provides a promising path toward understanding recessive genetic variation affecting early development.

## ACKNOWLEDGMENTS

This work was supported by NIH grants GM121828, GM089972, HG013015, and R35GM141906 to MVR, Charles H. Revson Foundation Fellowships to ABP and MVC, NIH fellowship GM090557 and NIH grant R35GM119744 to ABP, and NIH fellowship HD065442 to ASC. We gratefully acknowledge the *Caenorhabditis* Genetics Center, supported by NIH P40 OD010440. We thank Jasmine Nicodemus, Patrick Ammerman, Jia Shen, and Claire Curtin for help in the lab and members of the Rockman Lab for comments on drafts.

## SUPPLEMENTARY FILES

**Table S1. SibMatingExperiment.csv** Raw data reporting the results of the 1200 sibmating experiments described in the manuscript. For each experiment, the table reports the experiment number (used to blind the experimenters to the strain identity), an identifier for the worm picker who carried out the experiment, and the worm strain and species. The result for each experiment is recorded as the last generation of the experiment in which the strain was observed alive. The status column indicates whether the strain died in that last recorded generation (status = 1) or whether it was still alive (status = 0). The columns Copulation, LaidEmbryos, and Young Adults record for each experiment whether in the final generation the line successfully copulated, laid embryos, and produced offspring that developed to the Young Adult stage. The data in this file underlie Figure 1 and Table 1.

**Table S2. RegressionResults.csv** For each parameter (*A, B*) and each trait (Reproduction, Copulation, Fertility, Growth), the table reports the estimate, standard error, *t*-statistic, and p-value for each of the 11 isolates. For inbreeding load *B*, the table also shows p-values estimated from 10,000 simulations under the null hypothesis of no load.

**Table S3. RelativeFitnessData.csv** For each of 120 individual females tracked through their reproductive lives, the table reports the strain, species, experimental plate, and number of male and female adults that developed from each of eight 8-hour time blocks. For example, columns 1F and 1M record the Female and Male progeny that developed to adulthood from embryos laid during the 1^st^ time block. These data underlie Figure 2.

**Table S4. ScannerFitnessData.csv** For each of 59 experimental plates tracked by flatbed scanner for population growth, the table records the genotype (*treatment*), the phenotype (*sd_hours_to_starve*, hours to resource exhaustion, measured as the peak of the standard deviation of pixel intensity), the replicate plate for the treatment (*rep*, 1-10), the scanner used to score the plate (*scanner*, 5-8), and the position of the replicate plate on the scanner (*pos*, 1-9). These data underlie Figure 3.

**File S1. Comparison of Reproduction Probability to Morton et al.’s Survival Probability**.

**File S2. SibMatingAnalyses.R**. This file contains annotated R code that performs the reported analyses of the sib mating experiments and reproduces Figure 1 and Tables 1 and S2, starting with the raw data from Table S1.

**File S3. CbeceiFitnessAnalyses.R**. This file contains annotated R code that performs the reported analyses of *C. becei* relative fitness and reproduces Figure 2, starting with the raw data from Table S3.

## REFERENCES

Adams, P. E., A. B. Crist, E. M. Young, J. H. Willis, P. C. Phillips, and J. L. Fierst. 2022. Slow recovery from inbreeding depression generated by the complex genetic architecture of segregating deleterious mutations. Mol Biol Evol 39:msab330.

Baldi, C., S. Cho, and R. E. Ellis. 2009. Mutations in two independent pathways are sufficient to create hermaphroditic nematodes. Science 326:1002–1005.

Barriere, A., S. P. Yang, E. Pekarek, C. G. Thomas, E. S. Haag, and I. Ruvinsky. 2009. Detecting heterozygosity in shotgun genome assemblies: Lessons from obligately outcrossing nematodes. Genome Res 19:470–480.

Bataillon, T. and M. Kirkpatrick. 2000. Inbreeding depression due to mildly deleterious mutations in finite populations: Size does matter. Genet Res 75:75–81.

Ben-David, E., A. Burga, and L. Kruglyak. 2017. A maternal-effect selfish genetic element in Caenorhabditis elegans. Science 356:1051–1055.

Braendle, C. and A. Paaby. 2024. Life history in Caenorhabditis elegans: From molecular genetics to evolutionary ecology. Genetics 228:iyae151.

Brown, K. E. and J. K. Kelly. 2020. Severe inbreeding depression is predicted by the “rare allele load” in Mimulus guttatus. Evolution 74:587–596.

Charlesworth, D. and B. Charlesworth. 1987. Inbreeding depression and its evolutionary consequences. Annu Rev Ecol Syst 18:237–268.

Charlesworth, D. and J. H. Willis. 2009. Fundamental concepts in genetics the genetics of inbreeding depression. Nat Rev Genet 10:783–796.

Cutter, A. D. 2019. Reproductive transitions in plants and animals: Selfing syndrome, sexual selection and speciation. New Phytol 224:1080–1094.

Cutter, A. D., L. T. Morran, and P. C. Phillips. 2019. Males, outcrossing, and sexual selection in Caenorhabditis nematodes. Genetics 213:27–57.

Dey, A., C. K. Chan, C. G. Thomas, and A. D. Cutter. 2013. Molecular hyperdiversity defines populations of the nematode Caenorhabditis brenneri. Proc Natl Acad Sci USA 110:11056–11060.

Dolgin, E. S., B. Charlesworth, S. E. Baird, and A. D. Cutter. 2007. Inbreeding and outbreeding depression in Caenorhabditis nematodes. Evolution 61:1339–1352.

Dussex, N., H. E. Morales, C. Grossen, L. Dalen, and C. van Oosterhout. 2023. Purging and accumulation of genetic load in conservation. Trends Ecol Evol 38:961–969.

Ebel, E. R. and P. C. Phillips. 2016. Intrinsic differences between males and females determine sex-specific consequences of inbreeding. BMC Evol Biol 16:36.

Farhadifar, R., C. F. Baer, A. C. Valfort, E. C. Andersen, T. Muller-Reichert, M. Delattre, and D. J. Needleman. 2015. Scaling, selection, and evolutionary dynamics of the mitotic spindle. Curr Biol 25:732–740.

Farhadifar, R., C. H. Yu, G. Fabig, H. Y. Wu, D. B. Stein, M. Rockman, T. Muller-Reichert, M. J. Shelley, and D. J. Needleman. 2020. Stoichiometric interactions explain spindle dynamics and scaling across 100 million years of nematode evolution. Elife 9:e55877.

Felix, M. A., C. Braendle, and A. D. Cutter. 2014. A streamlined system for species diagnosis in Caenorhabditis (Nematoda: Rhabditidae) with name designations for 15 distinct biological species. PLoS One 9:e94723.

Fierst, J. L., J. H. Willis, C. G. Thomas, W. Wang, R. M. Reynolds, T. E. Ahearne, A. D. Cutter, and P. C. Phillips. 2015. Reproductive mode and the evolution of genome size and structure in Caenorhabditis nematodes. PLoS Genet 11:e1005323.

Glemin, S. 2003. How are deleterious mutations purged? Drift versus nonrandom mating. Evolution 57:2678–2687.

Hedrick, P. W. and A. Garcia-Dorado. 2016. Understanding inbreeding depression, purging, and genetic rescue. Trends Ecol Evol 31:940–952.

Kimura, M., T. Maruyama, and J. F. Crow. 1963. The mutation load in small populations. Genetics 48:1303–1312.

Kiontke, K. C., M. A. Felix, M. Ailion, M. V. Rockman, C. Braendle, J. B. Penigault, and D. H. Fitch. 2011. A phylogeny and molecular barcodes for Caenorhabditis, with numerous new species from rotting fruits. BMC Evol Biol 11:339.

Lande, R. and D. W. Schemske. 1985. The evolution of self-fertilization and inbreeding depression in plants. I. Genetic models. Evolution 39:24–40.

Levin, M., T. Hashimshony, F. Wagner, and I. Yanai. 2012. Developmental milestones punctuate gene expression in the caenorhabditis embryo. Dev Cell 22:1101–1108.

Lloyd, D. G. 1979. Some reproductive factors affecting the selection of self-fertilization in plants. Am Nat 113:67–79.

Lynch, M. and B. Walsh. 1998. Genetics and analysis of quantitative traits. Sinauer Associates, Sunderland, Massachusetts.

Memar, N., S. Schiemann, C. Hennig, D. Findeis, B. Conradt, and R. Schnabel. 2019. Twenty million years of evolution: The embryogenesis of four Caenorhabditis species are indistinguishable despite extensive genome divergence. Dev Biol 447:182–199.

Morton, N. E., J. F. Crow, and H. J. Muller. 1956. An estimate of the mutational damage in man from data on consanguineous marriages. Proc Natl Acad Sci USA 42:855–863.

Noble, L. M., A. S. Chang, D. McNelis, M. Kramer, M. Yen, J. P. Nicodemus, D. D. Riccardi, P. Ammerman, M. Phillips, T. Islam, and M. V. Rockman. 2015. Natural variation in plep-1 causes male-male copulatory behavior in C. elegans. Curr Biol 25:2730–2737.

Noble, L. M., J. Yuen, L. Stevens, N. Moya, R. Persaud, M. Moscatelli, J. L. Jackson, G. Zhang, R. Chitrakar, L. R. Baugh, C. Braendle, E. C. Andersen, H. S. Seidel, and M. V. Rockman. 2021. Selfing is the safest sex for Caenorhabditis tropicalis. Elife 10:e62587.

Paaby, A. B., A. G. White, D. D. Riccardi, K. C. Gunsalus, F. Piano, and M. V. Rockman. 2015. Wild worm embryogenesis harbors ubiquitous polygenic modifier variation. Elife 4:e09178.

Palopoli, M. F., M. V. Rockman, A. TinMaung, C. Ramsay, S. Curwen, A. Aduna, J. Laurita, and L. Kruglyak. 2008. Molecular basis of the copulatory plug polymorphism in Caenorhabditis elegans. Nature 454:1019–1022.

Robinson, J., C. C. Kyriazis, S. C. Yuan, and K. E. Lohmueller. 2023. Deleterious variation in natural populations and implications for conservation genetics. Annu Rev Anim Biosci 11:93–114.

Ross, J. A., D. C. Koboldt, J. E. Staisch, H. M. Chamberlin, B. P. Gupta, R. D. Miller, S. E. Baird, and E. S. Haag. 2011. Caenorhabditis briggsae recombinant inbred line genotypes reveal inter-strain incompatibility and the evolution of recombination. PLoS Genet 7:e1002174.

Salome-Correa, J.A., L. M. Noble, S. A. Sloat, T. H. M. Nguyen, and M. V. Rockman. 2025. Conservative evolution of genetic and genomic features in Caenorhabditis becei, an experimentally tractable gonochoristic worm. bioRxiv doi: 10.1101/2025.05.09.653148.

Schedl, T. and J. Kimble. 1988. fog-2, a germ-line-specific sex determination gene required for hermaphrodite spermatogenesis in Caenorhabditis elegans. Genetics 119:43–61.

Sin, S. Y. W., B. A. Hoover, G. A. Nevitt, and S. V. Edwards. 2021. Demographic history, not mating system, explains signatures of inbreeding and inbreeding depression in a large outbred population. Am Nat 197:658–676.

Sloat, S. A., L. M. Noble, A. B. Paaby, M. Bernstein, A. Chang, T. Kaur, J. Yuen, S. C. Tintori, J. L. Jackson, A. Martel, J. A. Salome Correa, L. Stevens, K. Kiontke, M. Blaxter, and M. V. Rockman. 2022. Caenorhabditis nematodes colonize ephemeral resource patches in neotropical forests. Ecol Evol 12:e9124.

Stein, L. D., Z. Bao, D. Blasiar, T. Blumenthal, M. R. Brent, N. Chen, A. Chinwalla, L. Clarke, C. Clee, A. Coghlan, A. Coulson, P. D’Eustachio, D. H. Fitch, L. A. Fulton, R. E. Fulton, S. Griffiths-Jones, T. W. Harris, L. W. Hillier, R. Kamath, P. E. Kuwabara, et al. 2003. The genome sequence of Caenorhabditis briggsae: A platform for comparative genomics. PLoS Biol 1:e45.

Sun, S., N. Kanzaki, M. Dayi, Y. Maeda, A. Yoshida, R. Tanaka, and T. Kikuchi. 2022. The compact genome of Caenorhabditis niphades n. sp., isolated from a wood-boring weevil, niphades variegatus. BMC Genomics 23:765.

Teterina, A. A., J. H. Willis, M. Lukac, R. Jovelin, A. D. Cutter, and P. C. Phillips. 2023. Genomic diversity landscapes in outcrossing and selfing Caenorhabditis nematodes. PLoS Genetics 19:e1010879.

Teterina, A. A., J. H. Willis, and P. C. Phillips. 2020. Chromosome-level assembly of the Caenorhabditis remanei genome reveals conserved patterns of nematode genome organization. Genetics 214:769–780.

Tintori, S. C., D. Çağlar, P. Ortiz, I. Chyzhevskyi, T. A. Mousseau, and M. V. Rockman. 2024. Environmental radiation exposure at Chornobyl has not systematically affected the genomes or chemical mutagen tolerance phenotypes of local worms. Proc Natl Acad Sci USA 121:e2314793121.

Tintori, S. C., D. Çağlar, and M. V. Rockman. 2025. A multigenerational population-growth assay to capture subtle fitness phenotypes in C. elegans and other nematodes. Genetics advance of press iyaf073.

Torres Cleuren, Y. N., C. K. Ewe, K. C. Chipman, E. R. Mears, C. G. Wood, C. E. A. AlAlami, M. R. Alcorn, T. L. Turner, P. M. Joshi, R. G. Snell, and J. H. Rothman. 2019. Extensive intraspecies cryptic variation in an ancient embryonic gene regulatory network. Elife 8:e48220.

Verster, A. J., A. K. Ramani, S. J. McKay, and A. G. Fraser. 2014. Comparative rnai screens in C. elegans and C. briggsae reveal the impact of developmental system drift on gene function. PLoS Genet 10:e1004077.

Wade, M. J. 1998. The evolutionary genetics of maternal effects. Maternal Effects as Adaptations, 5–21.

Wolf, J. B. and M. J. Wade. 2016. Evolutionary genetics of maternal effects. Evolution 70:827–839.

Yin, D., E. M. Schwarz, C. G. Thomas, R. L. Felde, I. F. Korf, A. D. Cutter, C. M. Schartner, E. J. Ralston, B. J. Meyer, and E. S. Haag. 2018. Rapid genome shrinkage in a self-fertile nematode reveals sperm competition proteins. Science 359:55–61.

Zhao, Z., T. J. Boyle, Z. Bao, J. I. Murray, B. Mericle, and R. H. Waterston. 2008. Comparative analysis of embryonic cell lineage between Caenorhabditis briggsae and Caenorhabditis elegans. Dev Biol 314:93–99.

