## Supplementary material for "Variation in inbreeding depression within and among *Caenorhabditis* species": FileS1.Model.pdf

**File S1:**
**Comparison of reproduction probability to Morton *et al.*'s survival probability**

Morton *et al.* model the probability of survival ($S$) given a single recessive locus as $S = 1 - Fqs - (1\text{-}F)q^2s - (1\text{-}F)2pqsh,$ where $s$ is the probability that an individual homozygous for the recessive allele fails to survive, $q$ and $p$ are the mutant allele frequency and its complement, $h$ is the dominance coefficient, and $F$ is the inbreeding coefficient. We substitute mating success for survival, and model the probability that two worms successfully mate, which requires that both worms succeed. Thus the probability of successful reproduction ($R$) is the product of the two worms' individual reproduction probabilities (i.e., $R = S^2$), yielding $R = 1 - 2Fqs + F^2q^2s^2 - 2(1\text{-}F)q^2s + 2F(1\text{-}F)q^3s^2 - 4(1\text{-}F)pqsh + 4F(1\text{-}F)pq^2s^2h + (1\text{-}F)^2q^4s^2 + 4(1\text{-}F)^2pq^3s^2h + 4(1\text{-}F)^2p^2q^2s^2h^2$.

In this case, unlike that of Morton *et al.,* the probability is quadratic rather than linear as a function of $F$. However, as shown below, all of the $F^2$ terms include $q^2s^2$ and are negligible.

Morton *et al.* extended their model to include the effects of multiple loci and environmental causes of individual failure ($x$). Assuming that these causes act independently, $S = \prod(1\text{-}x)(1 - Fqs - (1\text{-}F)q^2s - (1\text{-}F)2pqsh)$, where the product is over all loci and environmental causes. Using the approximation $1-t \sim e^{-t}$ for small $t,$ and assuming that environmental effects $x$ act independently on the two animals in each cross,
$R = \exp(-2\sum x + \sum x^2 - 2\sum q^2s - 4\sum pqsh + \sum q^4s^2 + 4\sum pq^3s^2h + 4\sum p^2q^2s^2h^2 - 2F\sum qs + 2F\sum q^2s + 2F\sum q^3s^2 + 4F\sum pqsh + 4F\sum pq^2s^2h - 2F\sum q^4s^2 - 8F\sum pq^3s^2h - 8F\sum p^2q^2s^2h^2 + F^2\sum q^2s^2 - 2F^2\sum q^3s^2 - 4F^2\sum pq^2s^2h + F^2\sum q^4s^2 + 4F^2\sum pq^3s^2h + 4F^2\sum p^2q^2s^2h^2)$

Thus,
$-\log R = A_R + B_RF + C_RF^2$
where $A_R = 2\sum x - \sum x^2 + \sum qs(2q + 4ph - q^2s - 4pq^2sh - 4p^2qsh^2)$
$B_R = 2\sum qs(1 - q - q^2s - 2ph - 2pqsh + q^3s + 4pq^2sh + 4p^2qsh^2)$
and $C_R = \sum q^2s^2(-1 + 2q + 4ph - q^2 - 4pqh - 4p^2h^2)$.

$B_R$ and $C_R$ are both zero when $h = 0.5$. The quadratic coefficient $C_R$ is much less than 1% of $B_R$ for any plausible values of *s, h,* and *q*.

The coefficients $A_R$ and $B_R$ here are approximately twice the magnitude of those in the single-individual case of Morton et al.
$B_{Survival} = \sum qs(1 - q - 2ph)$
$B_R = 2B_{Survival} + 2\sum q^2s^2(-q - 2ph + q^2 + 4pqh + 4ph^2)$

Similarly,
$A_{Survival} = \sum x + \sum qs(q + 2ph)$
$A_R = 2A_{Survival} - \sum x^2 - \sum q^2s^2(q + 4pqh + 4p^2h^2)$
