## Supplementary material for "Variation in inbreeding depression within and among *Caenorhabditis* species": FileS2.SibMatingAnalyses.R

```
# File S2. Analyses
# R version 4.3.3
# x86_64-apple-darwin20

#inbreeding coefficients for the parents of each cross, from recursion equation for full sibs, taken from Lynch and Walsh eq 10.5b. 
fs <- NULL;
fs[1:2] <- c(0,0)
for(i in 3:25){fs[i] <- 0.25*(1+ 2*fs[i-1]+fs[i-2])}

#Read in the inbreeding data
inb <- read.csv(file = "TableS1.SibMatingExperiment.csv", stringsAsFactors = T)

# Very inefficient loops for making a data.frame of strain x generation results
AllCrossFailures <- NULL;
AllCrossSuccesses <- NULL;
Allfstats <- NULL; 
AllStrains <- NULL;
AllCopSuccess <- NULL;
AllCopFailures <- NULL;
AllFertSuccess <- NULL;
AllFertFailures <- NULL;
AllGroSuccess <- NULL;
AllGroFailures <- NULL;

for(q in 1:11){ #For every strain...

	Failure <- NULL; 
	Success <- NULL;
	fstat <- NULL;
	strain <- NULL;
	cop <- NULL;
	Cod <- NULL; 
	fert <- NULL;
	Ste <- NULL;
	gro <- NULL;
	EmbLva <- NULL;

	straindata <- inb[inb$Strain == levels(inb$Strain)[q],]
	
		
		for(i in 0:20){	#For every generation...
			
			Failure[i+1] <- sum(straindata $LastGen == i & straindata $Status == 1, na.rm=T) #this right-censors the few plates that were lost during the experiment
			Success[i+1] <- sum(straindata $LastGen > i , na.rm = T)
			fstat[i+1] <- fs[i+1]; # this applies the parental generation f statistic to each cross. 
			strain[i+1] <- levels(inb$Strain)[q]
			cop[i+1] <- sum(straindata$LastGen ==i & straindata$Status ==1 & straindata$Copulation == 1, na.rm =T)
			Cod[i+1] <- sum(straindata$LastGen ==i & straindata$Status ==1 & straindata$Copulation == 0, na.rm =T)
			fert[i+1] <- sum(straindata$LastGen ==i & straindata$Status ==1 & straindata$LaidEmbryos == 1, na.rm = T)
			Ste[i+1] <- sum(straindata$LastGen ==i & straindata$Status ==1 & straindata$Copulation == 1 & straindata$LaidEmbryos == 0, na.rm = T)			
			gro[i+1] <- sum(straindata$LastGen ==i & straindata$Status ==1 & straindata$YoungAdults == 1 , na.rm = T);			
			EmbLva[i+1] <-  sum(straindata$LastGen ==i & straindata$Status ==1 & straindata$LaidEmbryos == 1 & straindata$YoungAdults == 0, na.rm = T)
		
		}
		
		###
		#This block combines the first and second cross generations, where F=0 for both. 
		F0failures = sum(Failure[fstat == 0])
		F0successes = sum(Success[fstat == 0])
		F0cop = sum(cop[fstat == 0])
		F0Cod =sum(Cod[fstat ==0])
		F0fert = sum(fert[fstat == 0]) 
		F0Ste = sum(Ste[fstat == 0])
		F0gro  = sum(gro[fstat ==0])
		F0Emb = sum(EmbLva[fstat == 0])
		Failure <- c(F0failures, Failure[3:21])
		Success <- c(F0successes, Success[3:21])
		cop <- c(F0cop, cop[3:21])
		Cod <- c(F0Cod, Cod[3:21])
		fert <- c(F0fert, fert[3:21])
		Ste <- c(F0Ste, Ste[3:21])
		gro <- c(F0gro, gro[3:21])
		EmbLva <-c(F0Emb, EmbLva[3:21])
		fstat <- fstat[2:21]
		strain <- strain[2:21]
		###
			
	# Concatenate data across strains
	AllCrossFailures <- c(AllCrossFailures, Failure)
	AllCrossSuccesses <- c(AllCrossSuccesses, Success)
	Allfstats <- c(Allfstats, fstat)
	AllStrains <- c(AllStrains, strain)
	AllCopSuccess <- c(AllCopSuccess, cop);
	AllCopFailures <- c(AllCopFailures, Cod)
	AllFertSuccess <- c(AllFertSuccess, fert);
	AllFertFailures <- c(AllFertFailures, Ste);
	AllGroSuccess <- c(AllGroSuccess, gro)
	AllGroFailures <- c(AllGroFailures, EmbLva);


}
AllCopSuccesses <- AllCopSuccess + AllCrossSuccesses
AllFertSuccesses <- AllFertSuccess + AllCrossSuccesses
AllGroSuccesses <- AllGroSuccess + AllCrossSuccesses

Species <- vector(length = length(AllStrains))
Species[AllStrains == "EM464"] <- "USremanei"
Species[AllStrains == "PB219"] <- "USremanei"
Species[AllStrains == "QG549"] <- "OKremanei"
Species[AllStrains == "QG548"] <- "OKremanei"
Species[AllStrains == "QG702"] <- "Cpanamensis"
Species[AllStrains == "QG703"] <- "Cpanamensis"
Species[AllStrains == "QG704"] <- "Cbecei"
Species[AllStrains == "QG711"] <- "Cbecei"
Species[AllStrains == "QG122"] <- "Ckamaaina"
Species[AllStrains == "QG123"] <- "Ckamaaina"
Species[AllStrains == "fog-2"] <- "Celegans"
Species <- as.factor(Species)


#Four relevant datasets:
Reproduction <- data.frame(Yes = AllCrossSuccesses, No = AllCrossFailures, Strain = as.factor(AllStrains), f = Allfstats, Species)
Copulation <- data.frame(Yes = AllCopSuccesses, No = AllCopFailures, Strain = as.factor(AllStrains), f = Allfstats, Species)
Fertility <- data.frame(Yes = AllFertSuccesses, No = AllFertFailures, Strain = as.factor(AllStrains), f = Allfstats, Species)
Growth <- data.frame(Yes = AllGroSuccesses, No = AllGroFailures, Strain = as.factor(AllStrains), f = Allfstats, Species)

#Clean up
rm(list = c("AllCopFailures","AllCopSuccess","AllCopSuccesses","AllCrossFailures","AllCrossSuccesses","AllFertFailures","AllFertSuccess","AllFertSuccesses","Allfstats","AllGroFailures","AllGroSuccess","AllGroSuccesses","AllStrains","Cod","cop","EmbLva","F0Cod","F0cop","F0Emb","F0failures","F0fert","F0gro","F0Ste","F0successes","Failure","fert","fs","fstat","gro","i","q","Species","Ste","strain","straindata","Success"))


#################
# Morton, a function to perform iteratively-weighted linear regression of ln(success/total) on inbreeding coefficient f. 
#################
Morton <- function(SP){ #for example, Reproduction, a four-column dataframe of yes, no, strain, f
	
	SP$Yes[SP$Yes == 0] <- NA
	frame <- NULL;
	na.omit(data.frame(SP$Yes, SP$No, SP$Strain, SP$f)) -> frame
	names(frame) <- c("yes", "no",  "Strain", "Fs")	
	n <- frame$yes + frame$no
	lnR <- log(frame$yes/n)
	frame <- data.frame(frame, n, lnR)
	yescount <- frame$yes # yescount is here to adjust starting weights in cases where otherwise we'd have to divide by 0.
	yescount[frame$no ==0] <- frame$yes[frame$no == 0]-1
	wts <- (frame$n) * (yescount/frame$n) / (1-yescount/frame$n) # From Morton et al. 1956. Starting point for weights
	m1 <- lm(-lnR ~ -1 + Strain/Fs, data = frame, weights = wts)
	P <- exp(-predict(m1)) # estimate binomial success probability from the model 
	P[which(P > 0.999)] <- 0.999 # This prevents infinite and negative weights
	wts <- frame$n * P / (1-P) # iteratively reweight using the estimated probabilities
	m <- list();
	niter <- 20
	for(q in 1:niter){
 		m[[q]] <- lm(-lnR ~ -1 + Strain/Fs, weights = wts, data = frame)
		P <- exp(-predict(m[[q]]))
		P[which(P > 0.999)] <- 0.999
 		wts <- frame$n * P / (1-P)
	}
	model <- m[[niter]]
	return(model)	
}

##Example usage:
summary(Morton(Reproduction))
write.csv(data.frame(summary(Morton(Reproduction))$coef,summary(Morton(Copulation))$coef, summary(Morton(Fertility))$coef, summary(Morton(Growth))$coef ), "MortonResults.csv")

#################
# sim.Morton, a function to simulate the null distribution of the strain-specific t-statistics under the null hypothesis of no inbreeding depression (B = 0), conditional on the observed sample sizes. 
# Specify the dataset to simulate as SP = for example, Reproduction, a four-column dataframe of yes, no, strain, f
#Returns a matrix of simulated t-statistics, n.strains x n.sims.
#################

sim.Morton <- function(SP, n.sims = 100, n.strains = 11){
	StrainSpecific.t <- matrix(nrow = n.strains, ncol = n.sims)
	for(i in 1:n.sims){
		frame <- NULL;
		na.omit(data.frame(SP$Yes, SP$No, SP$Strain, SP$f)) -> frame
		names(frame) <- c("yes", "no",  "Strain", "Fs")
		n <- frame$yes + frame$no
		frame <- data.frame(frame, n)
		est.prob <- sum(frame$no, na.rm =T)/sum(frame$n, na.rm =T)
		rbinom(length(frame$no), frame$n, est.prob) -> frame$no
		frame$yes <- frame$n - frame$no
		frame$yes[frame$yes == 0] <- NA
		na.omit(frame) -> frame
		lnR <- log(frame$yes/frame$n)
		frame <- data.frame(frame, lnR)
		yescount <- frame$yes # yescount is here to adjust starting weights in cases where otherwise we'd have to divide by 0.
		yescount[frame$no ==0] <- frame$yes[frame$no == 0]-1
		wts <- (frame$n) * (yescount/frame$n) / (1-yescount/frame$n) # From Morton et al. 1956. Starting point for weights
		m1 <- lm(-lnR ~ -1 + Strain/Fs, data = frame, weights = wts)
		P <- exp(-predict(m1)) # estimate binomial success probability from the model 
		P[which(P > 0.999)] <- 0.999 # This prevents infinite and negative weights
		wts <- frame$n * P / (1-P) # iteratively reweight using the estimated probabilities
		m <- list();
		niter <- 20
		for(q in 1:niter){
	 		m[[q]] <- lm(-lnR ~ -1 + Strain/Fs, weights = wts, data = frame)
			P <- exp(-predict(m[[q]]))
			P[which(P > 0.999)] <- 0.999
	 		wts <- frame$n * P / (1-P)
		}
	StrainSpecific.t[1:n.strains,i] <-  summary(m[[niter]])$coef[(n.strains+1):(2*n.strains),3]  
	}
	return(StrainSpecific.t)	
}

#################
# EstPvaluesForBFromSimulations, a function to estimate p-values for the hypothesis that B = 0 using the t-statistic distributions from simulations. 
#Returns a vector of p-values, one for each strain
#################

EstPvaluesForBFromSimulations <- function(SP, n.sims = 100, n.strains = 11){
	model <- Morton(SP)
	model.t <- summary(model)$coef[(n.strains+1):(2*n.strains),3]
	sim.t <- sim.Morton(SP, n.sims, n.strains)
	pBs <- NULL;
	for(j in 1:n.strains){
		pBs[j] <- sum(sim.t[j,] > model.t[j])/n.sims
	}
	return(pBs)
}

##Example usage:
date()
EstPvaluesForBFromSimulations(Reproduction, n.sims = 100000); date()
EstPvaluesForBFromSimulations(Copulation, n.sims = 100000); date()
EstPvaluesForBFromSimulations(Fertility, n.sims = 100000); date()
EstPvaluesForBFromSimulations(Growth, n.sims = 100000); date()

#################
#Lynch's J statistic
#Test of the null hypothesis that B = 0.
#J = abs(^B) / sqrt(2*phi*VG0/nlines)
#phi is a function of the experimental design, as in Lynch 1988 Fig. 4.
#VG0 is the additive genetic variance for the phenotype in the founding population, i.e., Var(genetic part of A)
	# The estimated total variance in each founding gonochoristic population ranges from 0.000 to 0.036 depending on the phenotype and population.
#J > 1 implies that B differs from zero by more than two standard errors
#The number of lines per strain is a key input, and because it varies across the experiment the min is most conservative.
#################
Jstat <- function(SP, phi = 2.3, VG0 = 0.01, method = c("min", "mean", "geometric", "harmonic")){
	SP$Yes[SP$Yes == 0] <- NA
	frame <- NULL;
	na.omit(data.frame(SP$Yes, SP$No, SP$Strain, SP$f)) -> frame
	model <- Morton(SP)
	n.strains <- length(levels(SP$Strain))
	modelB <- summary(model)$coef[(n.strains+1):(2*n.strains),1]
	nlines <-  NULL;
	for(i in 1:n.strains){  #What's the number of lines assayed per observation
		obs <- (frame$SP.Yes+frame$SP.No)[frame$SP.Strain == levels(frame$SP.Strain)[i]]
		if(method == "min"){
			nlines[i] <- min(obs)		
		}
		else if(method == "mean"){
			nlines[i] <- mean(obs)					
		}
		else if(method == "geometric"){
			nlines[i] <- prod(obs)^(1/length(obs))					
		}
		else if(method == "harmonic"){			
			nlines[i] <- 1/mean(1/obs)					
		}else {
			stop("what method?")
		}
	}
	abs(modelB) / sqrt(2*phi*VG0/nlines)
}

##Example usage
Jstat(Reproduction, method = "min")


#################
#Plotting functions
library(RColorBrewer)
library(survival)
colseq <- c("blue",1, "blue","brown","brown","darkgreen","darkgreen","orange","orange","purple","purple")

#################
#MortonPlot, a function to plot the data and model for each strain, one at a time
#################
MortonPlot <- function(SP, strain){
	straincount <- length(levels(SP$Strain))
	model <- Morton(SP)
	modelA <- summary(model)$coef[1:straincount,1]
	modelB <- summary(model)$coef[(straincount+1):(2*straincount),1]
	modelB.P <- summary(model)$coef[(straincount+1):(2*straincount),4]
	SP$Yes[SP$Yes == 0] <- NA
	frame <- NULL;
	na.omit(data.frame(SP$Yes, SP$No, SP$Strain, SP$f)) -> frame
	names(frame) <- c("yes", "no",  "Strain", "Fs")
	lnR <- log(frame$yes/(frame$yes + frame$no))
	plot(-lnR[frame$Strain == strain] ~ Fs[frame$Strain == strain], cex = 0.05*(yes+no)[frame$Strain == strain],data = frame, ylim = c(-0.05,0.6), xlim = c(0,1.05), xaxs = "i", yaxs = "i", main = paste(strain, "\np =", signif(modelB.P[which(levels(SP$Strain) == strain)], digits = 2), "B =", round(modelB[which(levels(SP$Strain) ==strain)], 2)), xlab = "F", ylab = "-log(Probability)")
#	points(lnM[frame$Strain == strain] ~ Fs[frame$Strain == strain], cex = 0.01*model$weights[frame$Strain == strain],data = frame, col  = 2)
	abline(modelA[which(levels(SP$Strain) == strain)], modelB[which(levels(SP$Strain) == strain)])
}
#################
#Plot all of the strains
par(mfrow = c(2, 6));
for(x in c(1,3,6,7,4,5,8:11,2)){MortonPlot(Reproduction, strain = levels(Reproduction $Strain)[x])}

##################
#PlotRegressions, to show all the slopes and intercepts at once
PlotRegressions <- function(SP, sds = "no"){
	straincount <- length(levels(SP$Strain))
	model <- Morton(SP)
	modelA <- summary(model)$coef[1:straincount,1]
	modelAsd <- summary(model)$coef[1:straincount,2]	
	modelB <- summary(model)$coef[(straincount+1):(2*straincount),1]
	modelBsd <- summary(model)$coef[(straincount+1):(2*straincount),2]
	plot(x = c(0,1), y = c(0,0.4), ty = "n", xaxs = "i", yaxs = "i", xlab = "F", ylab = "-logR")
	for(i in 1:11){
		abline(modelA[i],modelB[i], lwd = 2, col = colseq[i])
		if(sds == "yes"){
			abline(modelA[i]+modelAsd[i], modelB[i]+modelBsd[i], lwd = .5, col = rainbow(11)[i])
			abline(modelA[i]-modelAsd[i], modelB[i]-modelBsd[i], lwd = .5, col = rainbow(11)[i])
		}
	}
}
PlotRegressions(Reproduction)
PlotRegressions(Copulation)
PlotRegressions(Fertility)
PlotRegressions(Growth)

##################
# Plot Figure 1

# Plot survival curves
par(mfrow = c(1,3))
# Extinction curves
plot(survfit(Surv(inb$LastGen, inb$Status)~inb$Strain), col = colseq, lwd =2, ylab = "Proportion of Lines Surviving", xlab = "Generation", yaxs = "i")
# Expected curves absent inbreeding depression
plot(survfit(Surv(inb$LastGen, inb$Status)~inb$Strain), col = colseq, lwd =0, ylab = "Expected Proportion of Lines Surviving", xlab = "Generation", yaxs = "i")
model <- Morton(Reproduction)
straincount <- length(levels(Reproduction$Strain))
modelA <- summary(model)$coef[1:straincount,1]
t <- seq(0,22,1)
for(i in 1:11){
lines(1-pexp(t,modelA[i])~I(t), lwd =2, lty = 3, col = colseq[i])
}
# Regression slopes
PlotRegressions(Reproduction)
```
