## Supplementary material for "Variation in inbreeding depression within and among *Caenorhabditis* species": FileS3.RelativeFitnessAnalyses.R

```
# Analysis of Fitness Data from Dec 2022 experiment

data <- read.csv(file = "TableS3.RelativeFitnessData.csv", head = T, stringsAsFactors=T) 
data <- data[data$Include == 1,]

# Reorder the levels to make the plots look nicer
data$Species <- factor(data$Species, levels = c("C. remanei", "C. rem OK", "C. kamaaina", "C. becei", "C. panamensis"))

data$Strain <- factor(data$Strain, levels = c("EM464","QG1777","QG1870","QG1956","QG549", "QG1818","QG1873","QG1962", "QG123", "QG1816","QG1910", "QG704", "QG1808","QG1901","QG1971", "QG702","QG1819","QG1892","QG1984"))


# Specify the phenotyping times
intervals <- c(6, 8,8,8,8,8,8,8) #hours between worm transfers, starting with time of mating
times <- cumsum(intervals)/24  #Timepoints measured in days

colseq <- c("blue","darkgreen","brown","purple","orange")

lastcol <- dim(data)[2]

data[,I(lastcol+1)] <- NA

#Calculate relative fitness for each female, with the appropriate founder mean fitness set to 1.
for(i in 1:5){
	species <- levels(data$Species)[i]
	founder <- levels(factor(data$Strain[data$Species == species]))[1]
	print(founder)
	founderrows <- which(data[data$Species == species,"Strain"] == founder)
	LE <- function(w){
		abs(1 - mean(apply(data[data$Species == species,5:20][founderrows,], 1, function(x){sum(exp(-w*times) * x, na.rm = T)})))  #Lotka-Euler function 
		}
	lambda <- optimize(LE, c(0,100))$minimum   #Find lambda that makes outbred fitness closest to 1.

	# Use the estimated lambda to calculate relative fitness from reproductive schedule data
	get.w <- function(x){sum(exp(-lambda*times) * x, na.rm = T)} #Lotka-Euler function 
	rel.w <- apply(data[data$Species == species,5:20], 1, get.w)
	data[data$Species == species,I(lastcol+1)] <- rel.w

}

names(data)[22] <- "RelativeFitness"
data[,23]<- apply(data[,5:20], 1, sum, na.rm = T)
names(data)[23] <- "NumberOffspring"

# Number of adult offspring is correlated with relative fitness
cor.test(data$RelativeFitness, data$NumberOffspring)

# Test for heterogeneity among inbred lines from each founder
for(i in 1:5){
	species <- levels(data$Species)[i]
	founder <- levels(factor(data$Strain[data$Species == species]))[1]
	print(founder)
	founderrows <- which(data[data$Species == species,"Strain"] == founder)
	inbreds <- data[data$Species == species, ][-founderrows,]
	inbreds$Strain <- factor(inbreds$Strain)
	print(anova(lm(RelativeFitness~Strain, data = inbreds)))
}

colseq <- c(rep("blue",4), rep("darkgreen",4),rep("brown",3), rep("purple",4), rep("orange",4))
stripchart(data$RelativeFitness~data$Strain, vert = T, pch = 19, cex = 0.0, cex.axis =0.8, ylab ="Relative Fitness")
points(seq(1:length(levels(data$Strain))),aggregate(data$RelativeFitness~ data$Strain, FUN = mean)[,2], col = colseq, pch = 21, cex = 3, lwd=2 )
stripchart(data$RelativeFitness~data$Strain, vert = T, pch = 19, cex = 0.4, jitter = 0.05, method = "jitter", add = T)
abline(v = c(0.5,4.5,8.5,11.5,15.5,19.5),  lwd = 3)
abline(v = c(1.5,5.5,9.5,12.5,16.5))
abline(h=1)


colseq <- c(rep("blue",4), rep("darkgreen",4),rep("brown",3), rep("purple",4), rep("orange",4))
stripchart(data$NumberOffspring ~data$Strain, vert = T, pch = 19, cex = 0.0, , cex.axis =0.8, ylab ="Number of Offspring")
points(seq(1:length(levels(data$Strain))),aggregate(data$NumberOffspring ~ data$Strain, FUN = mean)[,2], col = colseq, pch = 21, cex = 3, lwd=2 )
stripchart(data$NumberOffspring ~data$Strain, vert = T, pch = 19, cex = 0.4, jitter = 0.05, method = "jitter", add = T)
abline(v = c(0.5,4.5,8.5,11.5,15.5,19.5),  lwd = 3)
abline(v = c(1.5,5.5,9.5,12.5,16.5))
abline(h=1)
```
